# A Scalable and Modular Automated Pipeline for Stitching of Large Electron Microscopy Datasets

**DOI:** 10.1101/2021.11.24.469932

**Authors:** Gayathri Mahalingam, Russel Torres, Daniel Kapner, Eric T. Trautman, Tim Fliss, Sharmishtaa Seshamani, Eric Perlman, Rob Young, Samuel Kinn, JoAnn Buchanan, Marc Takeno, Wenjing Yin, Daniel Bumbarger, Ryder P. Gwinn, Julie Nyhus, Ed Lein, Stephen Smith, Clay Reid, Khaled Khairy, Stephan Saalfeld, Forrest Collman, Nuno Macarico da Costa

**Affiliations:** Allen Institute for Brain Science, Seattle, 98109, USA; HHMI Janelia Research Campus, Ashburn, 20147, USA; St. Jude Children’s Research Hospital, Memphis, 38105, USA; Yikes LLC, Baltimore, MD, USA; Epilepsy Surgery and Functional Neurosurgery, Swedish Neuroscience Institute, Seattle, WA, US

## Abstract

Serial-section electron microscopy (ssEM) is the method of choice for studying macroscopic biological samples at extremely high resolution in three dimensions. In the nervous system, nanometer-scale images are necessary to reconstruct dense neural wiring diagrams in the brain, so called *connectomes*. In order to use this data, consisting of up to 10^8^ individual EM images, it must be assembled into a volume, requiring seamless 2D stitching from each physical section followed by 3D alignment of the stitched sections. The high throughput of ssEM necessitates 2D stitching to be done at the pace of imaging, which currently produces tens of terabytes per day. To achieve this, we present a modular volume assembly software pipeline *ASAP* (Assembly Stitching and Alignment Pipeline) that is scalable to datasets containing petabytes of data and parallelized to work in a distributed computational environment. The pipeline is built on top of the *Render* [18] services used in the volume assembly of the brain of adult *Drosophila melanogaster* [2]. It achieves high throughput by operating on the meta-data and transformations of each image stored in a database, thus eliminating the need to render intermediate output. ASAP is modular, allowing for easy incorporation of new algorithms without significant changes in the workflow. The entire software pipeline includes a complete set of tools for stitching, automated quality control, 3D section alignment, and final rendering of the assembled volume to disk. ASAP has been deployed for continuous processing of several large-scale datasets of the mouse visual cortex and human brain samples including one cubic millimeter of mouse visual cortex [1, 25] at speeds that exceed imaging. The pipeline also has multi-channel processing capabilities and can be applied to fluorescence and multi-modal datasets like array tomography.

## Introduction

Serial section electron microscopy (ssEM) provides the high spatial resolution in the range of a few nanometers per pixel that is necessary to reconstruct the structure of neurons and their connectivity. However, imaging at a high resolution produces a massive amount of image data even for a volume that spans a few millimeters. For example, a cubic millimeter of cortical tissue imaged at a resolution of 4 × 4 × 40 nm^3^, generates more than a petabyte of data and contains more than 100 million individual image tiles [1]. These millions of images are then stitched in 2D for each section and aligned in 3D to assemble a volume that is then used for neuronal reconstruction. With parallelized high throughput microscopes producing tens of terabytes of data per day, it is necessary that this volume assembly process is automated and streamlined into a pipeline, so that it does not become a bottleneck. The ideal pipeline should be capable of processing data at the speed of imaging [30] and produce a high fidelity assembled volume. To match the speed of the EM imaging, the volume assembly pipeline needs to be automated to handle millions of images per day from multiple microscopes.

Several tools used in various stages of volume assembly pipelines perform image registration by extracting and matching similar features across overlapping images [6, 28, 8, 3]. Image registration using Fourier transformation [28] was used to successfully align mouse and zebrafish brain datasets acquired using wager mapper ssEM imaging technology. The Fiji [9, 10] plugin TrakEM2 [6] includes a comprehensive set of tools and algorithms to perform stitching and alignment of various types of microscopy image formats. AlignTK [8] implements scalable deformable 2D stitching and serial section alignment for large serial section datasets using local cross-correlation. An end-to-end pipeline to perform volume assembly and segmentation using existing tools were developed by R. Vescovi et al. [11] and was designed to run on varied computational systems. The pipeline was shown to process smaller datasets through supercomputers efficiently. While these approaches have been successfully used in the volume assembly of smaller datasets, they do not scale well for large scale datasets, lack support for different classes of geometric transformations, or do not incorporate reliable filters for false matches due to imaging artifacts [12].

We propose a volume assembly pipeline - *ASAP* (Assembly Stitching and Alignment Pipeline) that is capable of processing peta-scale EM datasets with high fidelity and processing rates that match the speed of imaging. Our pipeline is based on the volume assembly framework proposed in [2] and is capable of achieving high throughput by means of meta-data operations on every image in the dataset. The meta-data and transformations associated with each image are stored in a MongoDB database fronted by *Render* [18] services to dynamically render the output at any stage in the pipeline. The effectiveness of the pipeline has been demonstrated in the volume assembly of multiple peta-scale volumes.

The pipeline described here for assembly of large connectomics volumes is divided into two sections: 1) A software package that is scalable, modular, and parallelized and is deployable in varied computing environments to perform volume assembly of EM serial sections; 2) A workflow engine and a volume assembly workflow that utilizes these tools to automate the processing of raw EM images from a multi-scope setup using high performance computing (HPC) systems. The tools in ASAP are open-source and include abstract level functionalities to execute macro level operations in the pipeline. The modularity of the tools allows for easy implementation of other algorithms into the pipeline without making major changes to the existing setup. The software tools can be easily deployed in different computing environments such as HPC systems, cloud based services, or on a desktop computer in a production level setting. The software stack also includes a set of quality control tools that can be run in an automated fashion to assess the quality of the stitched montages. These software tools can be easily utilized by workflow managers running the volume assembly workflow to achieve high throughput. The tools are designed to generalize well for other datasets from different domains (that carry the assumption of generating overlapping images) and can be adapted to process such datasets. We have also developed a workflow manager *BlueSky* (github.com/AllenInstitute/blue_sky_workflow_engine) that implements the volume assembly workflow using our software stack. The proposed pipeline combined with *BlueSky* has been successfully used to stitch and align several high-resolution *mm*^3^ EM volume from the mouse visual cortex and a human dataset at speeds higher than the imaging rate of these serial sections from a highly parallelized multi-scope setup.

## Results

### Development of a stitching and alignment pipeline

The pipeline (ASAP) described in this work is based on the principles described by Kaynig et al. [4], Saalfeld et al. [5] and Zheng et al. [2], and scales the software infrastructure to stitch and align petascale datasets. It includes the following stages: 1) Lens distortion computation, 2) 2D Stitching 3) Global section-based nonlinear 3D alignment, 4) Fine 3D alignment and 5) Volume assembly. ASAP performs feature-based stitching and alignment in which point correspondences between two overlapping images are extracted and a geometric transformation is computed using these point correspondences to align the images.

Figure 1 shows the volume assembly pipeline (ASAP) for building 3D reconstruction out of serial section Transmission Electron Microscopy (ssTEM) images. First single images from serial sections from ssTEM are collected. As the field of view is limited, multiple images that overlap with each other are imaged to cover the entire section. Images acquired by ssTEMs can include dynamic nonlinear distortions brought about by the lens system. A compensating 2D Thin Plate Spline transformation is derived using a custom triangular mesh-based strategy [15] based on point correspondences of overlapping image tiles as in Kaynig et al. [4]. The point correspondences (also referred as point-matches) are extracted using SIFT [19] and a robust geometric consistency filter using a local optimization variant of RANSAC [21] and robust regression [5] (see Methods for more details). These point correspondences, in lens-corrected coordinates, are then used to find a per image affine/polynomial transformation that aligns the images in a section with each other to create a *montage*. The affine/polynomial transformations are computed using a custom Python package, BigFeta [7], which implements a direct global sparse matrix solving strategy based on Khairy et al. [12]. The stitched montages are then globally aligned with each other in 3D. The 3D global alignment is performed by extracting point correspondences between low-resolution version of the 2D stitched sections and solved with BigFeta to obtain a Thin Plate Spline per section transformation. This 3D alignment is the result of a progressive sequence of Rotational, Affine, and Thin Plate Spline solves with tuned regularization parameters such that each solution initializes the next more deformable, yet increasingly regularized transformation. The globally aligned transformations can then be used as an initialization for computing finer and denser alignment transformations (An example of this is the fine alignment described in [17]), which is computed on a per image basis at a much higher resolution. Several iterations of the global 3D alignment are performed to achieve a good initialization for the fine alignment process. For all the datasets presented in this manuscript, after global alignment the data was then transferred and fine alignment using SEAMLESS [17]) was applied.

**Figure 1:**
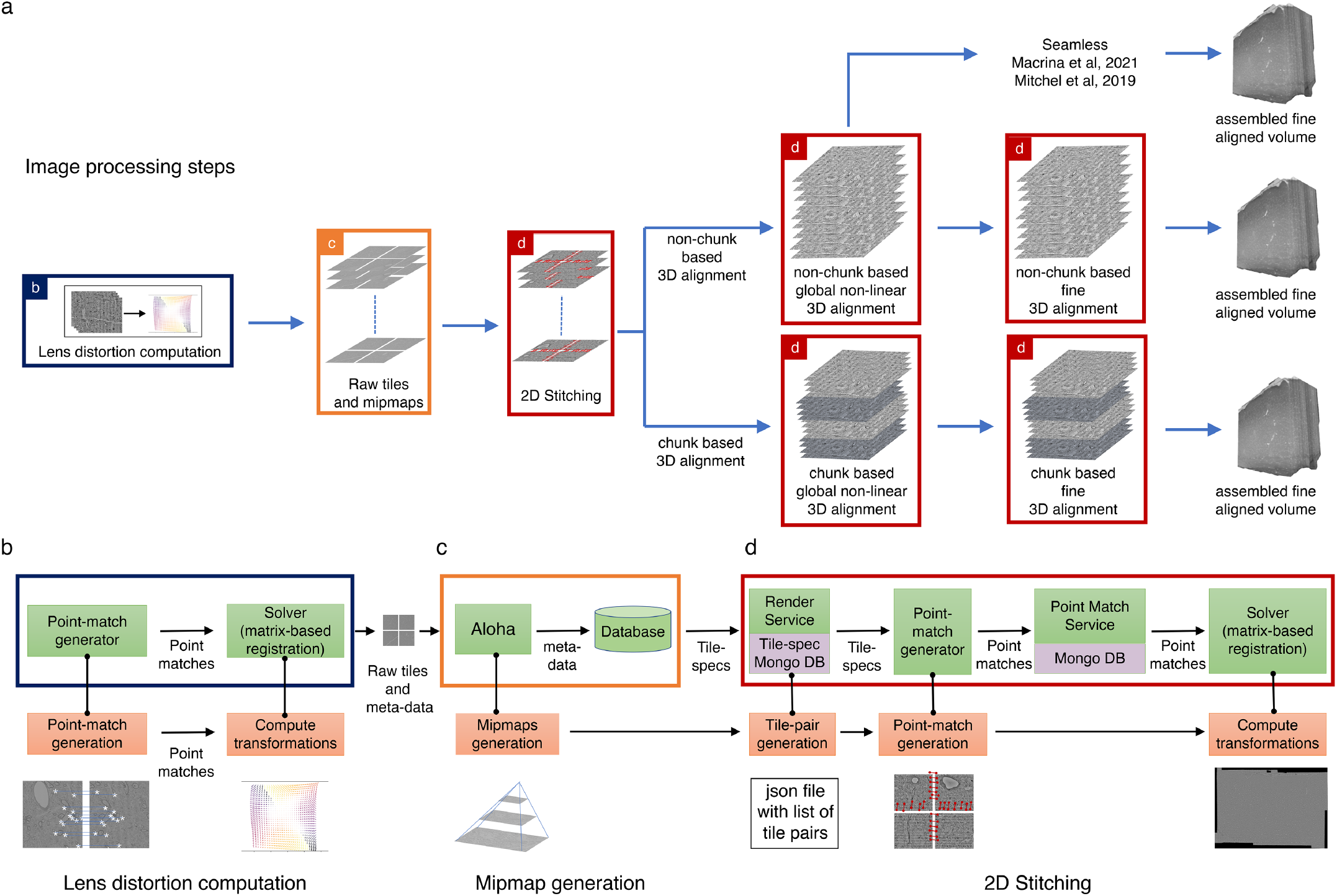
ASAP - Volume Assembly Workflow. (a) The different steps of image processing in *ASAP* for EM serial sections. The infrastructure permits multiple possible strategies for 3D alignment, including a chunk based approach in case it is not possible to 3D align the complete dataset at once, as well as using other workflows [17] (https://www.microns-explorer.org/cortical-mm3) for fine 3D alignment with the global 3D aligned volume obtained using ASAP. (b, c and d) Representation of different modules in the software infrastructure. The green boxes represent software components, the orange boxes represent processes and the purple processes represent databases. The color of the outline of the box matches its representation in the image processing steps shown in “a”. (b) Schematic showing the lens distortion computation. (c) Schematic describing the process of data transfer and storage along with MIPmaps generation using the data transfer service *Aloha*. (d) Schematic illustrating the montaging process of serial sections. The same software infrastructure of (d) is then also used for 3D alignment as shown by the red boxes in “a”.

In a continuous processing workflow scenario, the serial sections from multiple ssTEMs are stitched immediately once they are imaged. 3D alignment is performed on chunks of contiguous sections that partially overlap with their neighboring chunks. These independently 3D aligned chunks can be assembled to a full volume by aligning them rigidly and interpolating the transformations in the overlapping region (Figure 1).

### Software infrastructure supporting stitching and alignment

Our software infrastructure is designed to support EM imaging pipelines such as piTEAM [1] that produce multiple serial sections from a parallelized scope setup every hour. The infrastructure is designed for processing peta-scale datasets consisting of millions of partially overlapping EM images. The infrastructure consists of four core components: 1) A modular set of software tools that implements each stage of ASAP (asap-modules), 2) A service with REST APIs to transfer data from the microscopes to storage hardware (Aloha); 3) REST APIs for creating, accessing and modifying image meta-data (Render) and; 4) A matrix based registration system (BigFeta). Below we provide a brief description of these components with a more detailed description in the methods section ASAP modules.

ASAP is implemented as a modular set of tools that includes abstract level functions to execute for each stage of the volume assembly pipeline. It also includes quality control (QC) tools to assess stitching quality, to render results to disk at any stage of the pipeline, to obtain optimal parameters for computing point-correspondences, and to obtain optimal parameters for solving for optimal transformations. asap-modules is supported by *render-python* for read/writes to the database and *argschema* for its input and output data validation (See Methods section for more details).

Aloha is an image transfer service (Figure 1c) that receives raw images and their meta-data from the microscopes, stores them in primary data storage and losslessly compresses the original data to reduce the storage footprint. It includes REST APIs for clients to GET/POST images and their meta-data. It also produces downsampled representations of the images for faster processing and visualization.

Render [18] provides logic for image transformation, interpolation and rendering. It is backed by a MongoDB document store that contains JSON tile specifications with image meta-data and transformations. Render’s REST APIs are accessed by asap-modules using render-python to create, access and modify image metadata in the database. The REST APIs allow the user to access the current state of any given set of image tiles during the stitching process. Render also includes a point-match service that handles the storage and retrieval of point correspondences in a database, since computing point correspondences between millions of pairs of images is computationally expensive. Another advantage of storing the point correspondences in a database is that it is agnostic to the algorithm that is used for the computation of these point correspondences. The *point-match service* (Fig. 1c and e) handles the data ingestion and retrieval from the database using REST APIs with both operations being potentially massively distributed.

BigFeta [7] is a matrix-based registration system that estimates the image transformations using the point correspondences associated with the image. BigFeta includes transformations such as rotations to implement rigid alignments, and 2D Thin Plate Spline transformations that are useful for 3D image alignments. BigFeta can also be integrated with distributed solver packages such as PETSc [22] for solving large sparse matrices involving billions of point correspondences.

We also developed a workflow manager *BlueSky* as well as an associated volume assembly workflow to automatically process serial sections as they are continuously ingested during the imaging process. It utilizes the abstract level functions in asap-modules to create workflows for each stage of the volume assembly pipeline.

Our alignment pipelines operate only on meta-data (point correspondences and transformations) derived from image tiles - a feature derived from the Render services, thus allowing efficient processing of peta-scale datasets and the feasibility of real-time stitching with proper infrastructure. Where possible, the pipeline works with down-scaled versions of image tiles (MIPmaps) which dramatically increases processing speed and reduces disk usage as raw data can be moved to a long-term storage for later retrieval.

Beyond the use of this software infrastructure for EM data, which drove the development that we describe in this manuscript, the pipeline also has multi-channel processing capabilities and can be applied to fluorescence and multi-modal datasets like array tomography (see Figure 7)

### Data Acquisition and initiation of image processing

An important first step in our pipeline is the correction of lens distortion effects on raw images. Lens distortions are calculated from a special set of images with high tile overlap. These *calibration montages* are collected at least daily and after any event that might affect the stability of the beam (e.g. filament replacement). This step is followed by the acquisition of the neuroanatomical dataset, for which a bounding box is drawn around the ROI in each ultra-thin section. In certain situations, multiple ROIs are required per section. The volume assembly workflow accepts multiple entries referencing the same placeholder label to support re-imaging. At the end of each acquisition session, the tiles, tile manifest, and session log are uploaded to the data center storage cluster and the lens correction and montaging workflows in the volume assembly workflow are triggered. Figure 2 shows the specialized services that facilitate data transfer and tracking from high-throughput microscopes to shared compute resources.

**Figure 2:**
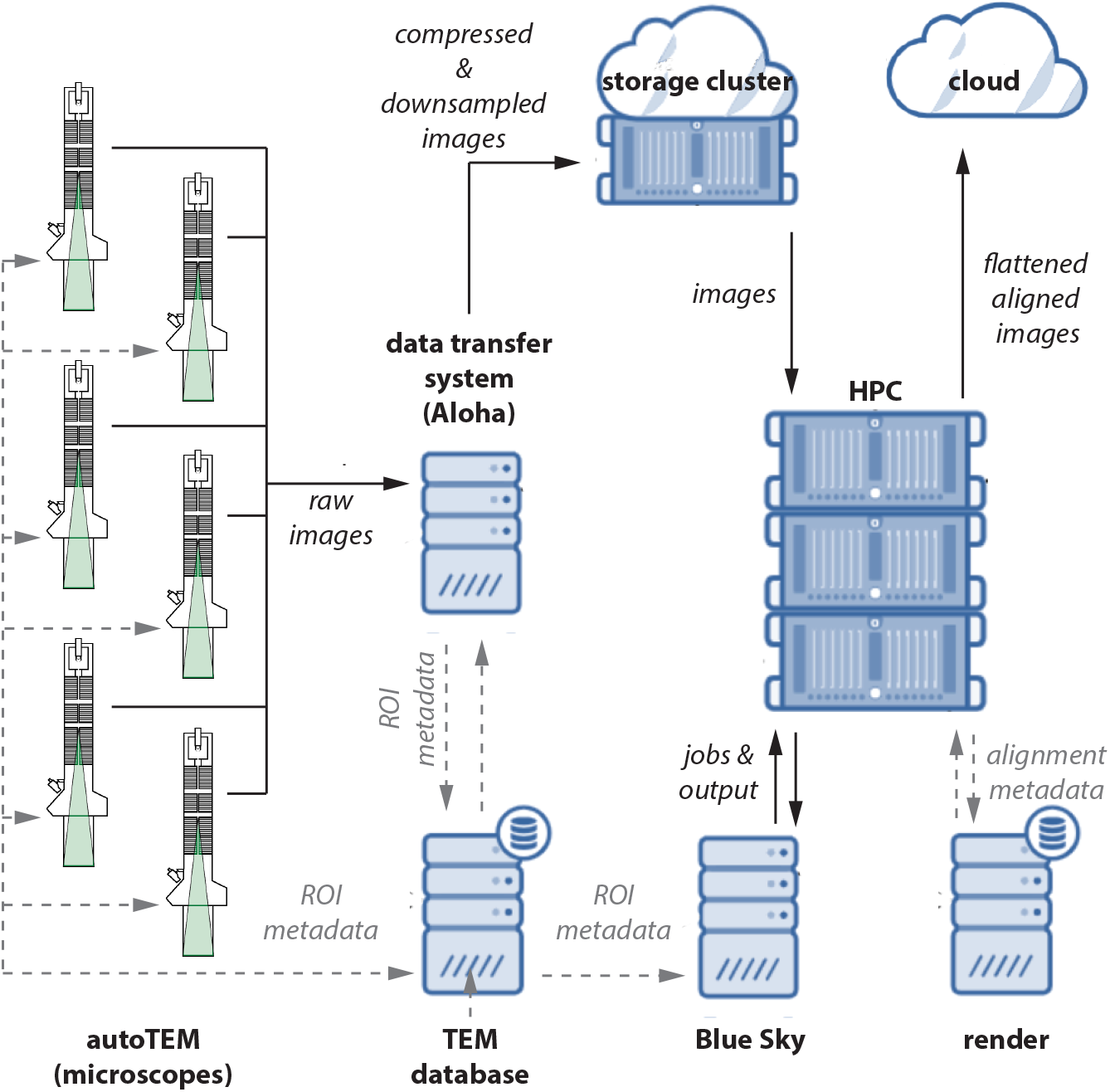
Data flow diagram. A schematic diagram showing the flow of image data, meta-data, and processed data between microscopes. Raw images and meta-data are transferred from microscopes to our data transfer system *(Aloha)* and TEM database, respectively. Aloha generates MIPmaps and compresses images and transfers them to the storage cluster for further processing by *ASAP*. Meta-data is transferred to BlueSky through TEM database, which triggers the stitching and alignment process. The meta-data from the stitching process is saved in the Render services database. The final assembled volume is transferred to the cloud for further fine alignment and segmentation.

This infrastructure was used to process multiple petascale datasets, including a 1 mm^3^ (mouse dataset 1) of the mouse brain that is publicly available on microns-explorer.com [25]. Over 26,500 sections were imaged at 4nm/pixel resolution using 5 microscopes, running in a continuous and automated fashion [1]. Each montage is composed of ~5,000 tiles of 15 μm × 15 μm with an overlap of 13% in both *x* and *y* directions. The total file size of a single montage is about 80 GB and thus a daily throughput of 3.6 TB per system is produced in a continuous imaging scenario. Part of the dataset was imaged using a 50 MP camera with an increased tile size to 5,408 × 5,408 pixels. This resulted in montages with approximately 2,600 tiles at an overlap of 9% in both *x* and *y* directions. The infrastructure was also used to process two other large mouse datasets and a human dataset. The details about these datasets are shown in Table 1.

**Table 1:** Details of datasets processed using ASAP - Volume assembly pipeline

| Dataset | # of sections | Pixel resolution (nm/pixel) | Image size | Tile overlap | Total size (PiB) | Total size (non-overlap) (PiB) | ROI size (in microns) |
| --- | --- | --- | --- | --- | --- | --- | --- |
| Mouse dataset 1 | 19945 | 3.95-4 | 3840 × 3840 (20MP camera),<br>5408 × 5408 (50MP camera) | 13%, 9% | 1.6 | 1.2 | 1300 × 870 |
| Mouse dataset 2 | 17593 | 4.67-5 (4.78 mean) | 5376 × 5376 | 9-10%, 9.4% mean | 0.984 | 0.807 | 1191 × 815 |
| Mouse dataset 3 | 17310 | 3.8-4.15 (3.91 mean) | 5376 × 5376 | 9-10%, 9.1% mean | 0.665 | 0.549 | 647 × 672 |
| Human | 9673 | 4.65-4.95 (4.78 mean) | 5376 × 5376 | 9-10%, 9.5% mean | 1.18 | 0.968 | 2315 × 987 |

### Automated petascale stitching

Besides stitching and aligning large scale datasets, a requirement for the volume assembly pipeline is to achieve high fidelity at a rate that matches or exceeds the imaging speed, so as to provide rapid feedback on issues with the raw data encountered during the stitching process. This is achieved in our pipeline using an automated workflow manager (BlueSky) that executes the volume assembly pipeline to continuously process serial sections from 5 different autoTEMs [1].

The images from the autoTEMs are transferred to the Aloha service without sending them to storage servers directly. The Aloha service generates MIPmaps, compresses the raw images and then writes them to the storage servers. The sections processed by Aloha are then POSTed to the BlueSky workflow manager which initiates the montaging process. During an imaging run each microscope uploads raw data and meta-data to Aloha using a concurrent upload client. Limitations of the autoTEM acquisition computers cap the aloha client throughput at 0.8–1.2Gbps per microscope, which is sufficient for daily imaging with a 50 Megapixel camera as described in Yin et al. [1]. Transferring previously-imaged directories from high-performance storage servers has shown that an Aloha deployment on multiple machines is capable of saturating a 10 Gbps network uplink. The serial sections are assigned *pseudo z* indices to account for errors in meta-data from the scopes such as barcode reading errors that assigns incorrect z indices. The lens correction workflow is triggered to compute a transformation that can correct lens distortion effects on the raw images. This transformation is updated in the image meta-data so as to be used in subsequent stages of volume assembly. The montaging workflow in BlueSky triggers the generation of point correspondences and stores them in the database using the point-match service, followed by calculating the globally optimal affine/polynomial transformation for each image tile in the montage using the BigFeta solver. The transformations are saved as meta-data associated with each tile image in the Render services database. The montages go through an automated quality control (QC) process to ensure a high fidelity stitching (see Section Automated montage QC), followed by a global 3D alignment of the entire dataset.

ASAP is capable of performing the global 3D alignment in chunks, making it scalable to use in larger datasets or with fewer computational resources. However all our datasets have been 3D aligned as a single chunk. The montages are rendered to disk at a scale of 0.01 and point correspondences are computed between the neighboring sections represented by their downsampled versions. A per section Thin Plate Spline transformation is computed using 25-49 control points in a rectangular grid. The per-section transformation is then applied to all the tile images in that section to globally align them in 3D.

### Automated montage QC

Quality control is a crucial step at each stage of processing in EM volume assembly to ensure that the outcome at each stage is of high quality. ASAP-modules include a comprehensive set of tools to perform quality control of the computed lens correction transformations, the stitched montages, and the 3D aligned volume. These tools are integrated within the lens correction and montaging workflow in the volume assembly workflow to automatically compute statistical metrics indicating the stitching quality and also generates maps of montages showing potential stitching issues (see Figure 3). The stitched montages that pass QC are automatically moved to the next stage of processing thus enabling faster processing with minimal human intervention but ensuring a high quality volume assembly.

**Figure 3:**
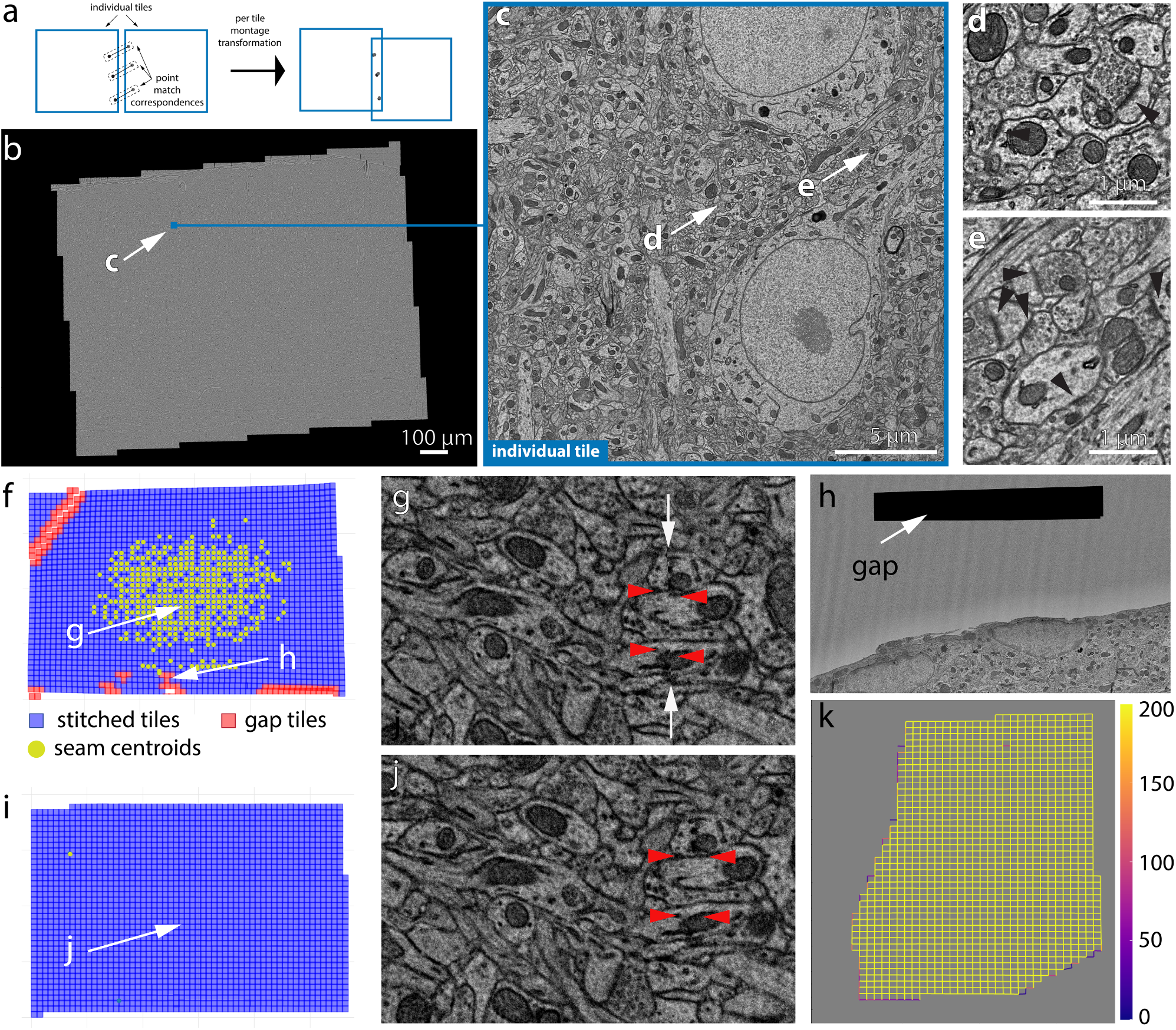
2D Stitching and automated assessment of montage quality. (a) Schematic diagram of the montage transformation using point correspondences. (b) Montage 2D stitched section from mouse dataset 1 (publicly available at www.microns-explorer.org [25]). (c) Single acquisition tile from the section in (b). (d and e) Detail of synapses (arrow heads) from the tile shown in (c). (f) QC plot of a stitched EM serial section with non-optimal parameters. Each blue square represent a tile image on how they appear aligned in the montage. The red squares represent tile images that have gaps in stitching with neighboring tile images and are usually located in regions with resin or film. (g) A zoom in region of the 2D montage in (f) showing the seam (white arrows) between tiles causing misalignment (red arrowheads) between membranes. (h) A zoomed in region of the section showing a tile having a gap with its neighbors. (i) QC plot of a stitched EM serial section after parameter optimization. (j) A zoom in region of the 2D montage in (i) showing no seams in the same region as in (g). The read arrow heads show the same locations as in (g). (k) A schematic plot representing the number of point correspondences between every pair of tile images for a section of the human dataset. Each edge of the squares in the plot represent the existence of point correspondences between tile images centered at the end points of the edge. The color of the edge represents the number of point correspondences computed between those tile image pairs.

The quality control maps (Fig. 3f and i) of the montages provide a rapid means to visually inspect and identify stitching issues associated with the montage without the need to materialize or visualize large scale serial sections. The QC map reveal the location of gaps and seams between tiles in addition to providing an accurate thumbnail representation of the stitched section. A seam (Fig. 3g) is defined as a misalignment between two tiles and is identified by means of the pixel residuals between point correspondences between the tiles. Misalignments can be eliminated by solving for correct transformations using optimized sets of parameters. A gap between tile images (Fig. 3h) is usually the result of inaccurate montage transformations that are caused by lack of point correspondences between tile pairs where the gap appears. Tile pairs that include features like blood vessel, resin or film region, etc. (see Fig. 3h) lack point correspondences thus causing a gap between the tiles during stitching. The stitching issues associated with the resin or film region are ignored, while the gaps in tiles containing blood vessels are solved with optimal parameters to ensure no misalignments between the tile and its neighbors. The density plot of point correspondences between tiles within a montage are shown in Fig. 3k and is a quick way to visualize areas with poor point correspondences, if any.

Sections that failed QC are examined by a human proofreader and moved to the appropriate stage of re-processing. Sections with insufficient point correspondences are sent to the point-matching stage of the montage workflow for generation of point correspondences at a higher resolution. Sections with misalignments are sent to the solver stage with new parameters. These parameters were heuristically chosen by means of a parameter optimization algorithm based on the stitching quality metrics (see Methods section Montage Parameter Optimization for more details).

Un-optimized parameters can also lead to distorted montages where individual tiles are distorted (see Figure 4c and d for distorted and un-distorted versions of the same montage). The Median Absolute Deviation (MAD) (Figure 4a and b) statistic provides a computational assessment of the quality of the montage and aids in the selection of optimized set of parameters to solve for a montage with high fidelity. The optimal *x* and *y* MAD statistic values were heuristically selected for every dataset.

**Figure 4:**
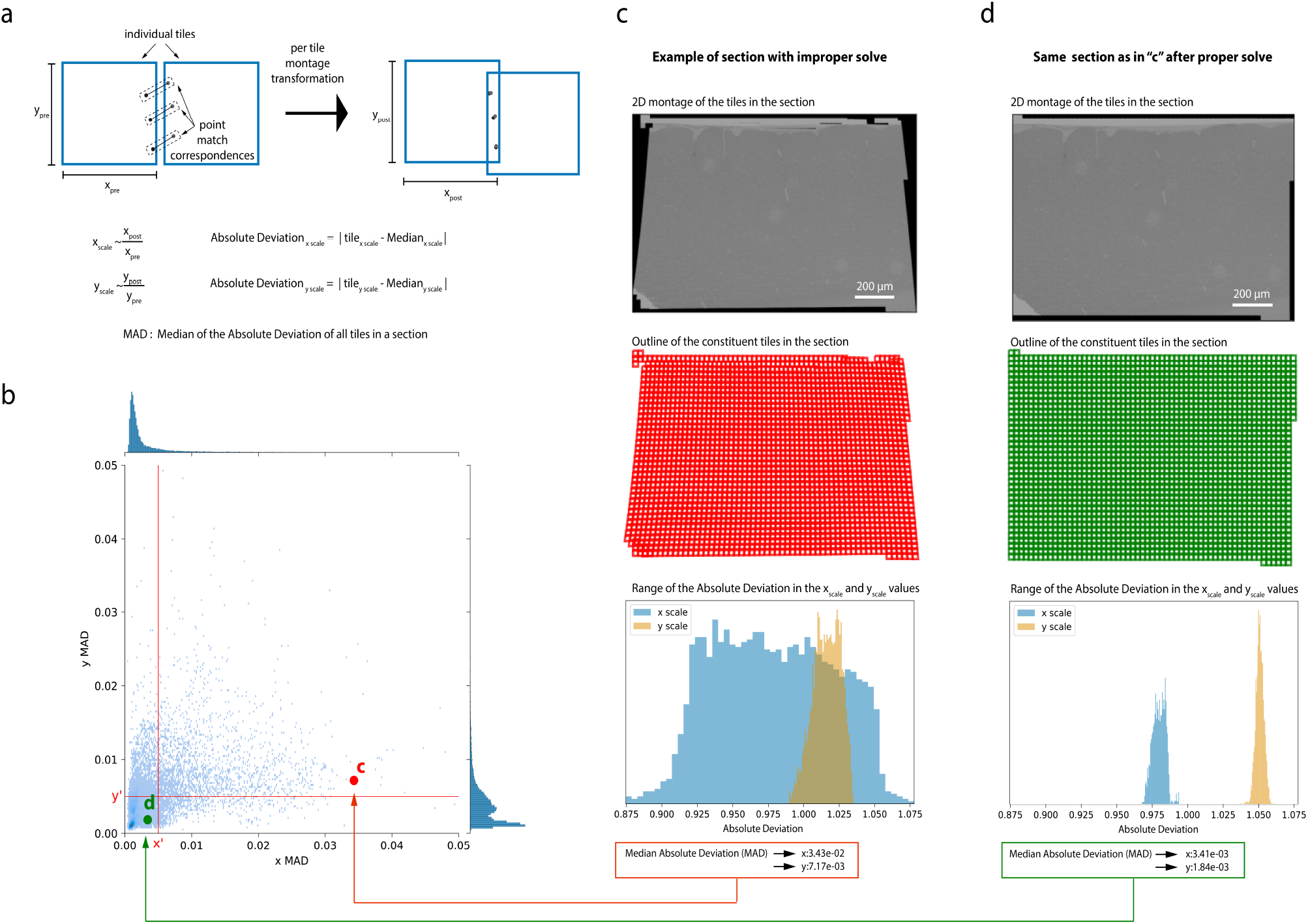
Median absolute deviation (MAD) statistics for montage distortion detection. (a) Schematic description of computation of Median Absolute Deviation (MAD) statistics for a montage. (b) A scatter plot of *x* and *y* MAD values for each montage. A good stitched section without distorted tile images falls in the third quadrant (where point (d) is shown). (c) An example of a distorted montage of a section solved using un-optimized set of parameters. Row 1 shows the downsampled version of the montaged section, row 2 shows the QC plot of the section showing the distortions, and row 3 shows the *x* and *y* absolute deviation distribution for the un-optimized montage. (d) Section shown in (c) solved with optimized parameters with row 1 showing the downsampled montage, row 2 showing the QC plot of the section, and row 3 showing the *x* and *y* absolute deviation distribution for the section.

### Performance of the volume assembly pipeline - ASAP

High quality 2D stitching and 3D alignment are necessary for accurate neuro-anatomy reconstruction and detection of synaptic contacts. The 2D stitching quality is assessed by a residual metric, which computes the sum of squared distances between point correspondences post stitching (see Fig 5a). A median residual of less than 5 pixels was achieved for sections from all our datasets (Fig. 5), which is a requirement for successful 3D segmentation ([17]) in addition to having no other stitching issues as described above. A small number of sections reported high residuals even with the optimized set of solver parameters (Fig. 5). An attempt to re-montage them with parameters that will reduce the residuals resulted in distorting individual tile images. Hence, these sections were montaged using a set of parameters that produces a montage with less distorted tiles and a residual that can be tolerated by the 3D fine alignment process and further segmentation. Overall, we aim to achieve high fidelity stitching by attempting to keep the residuals within the threshold, while preserving the image scales in both *x* and *y* closer to 1 (Fig. 5) and occasionally allowing montages with residuals above the threshold.

**Figure 5:**
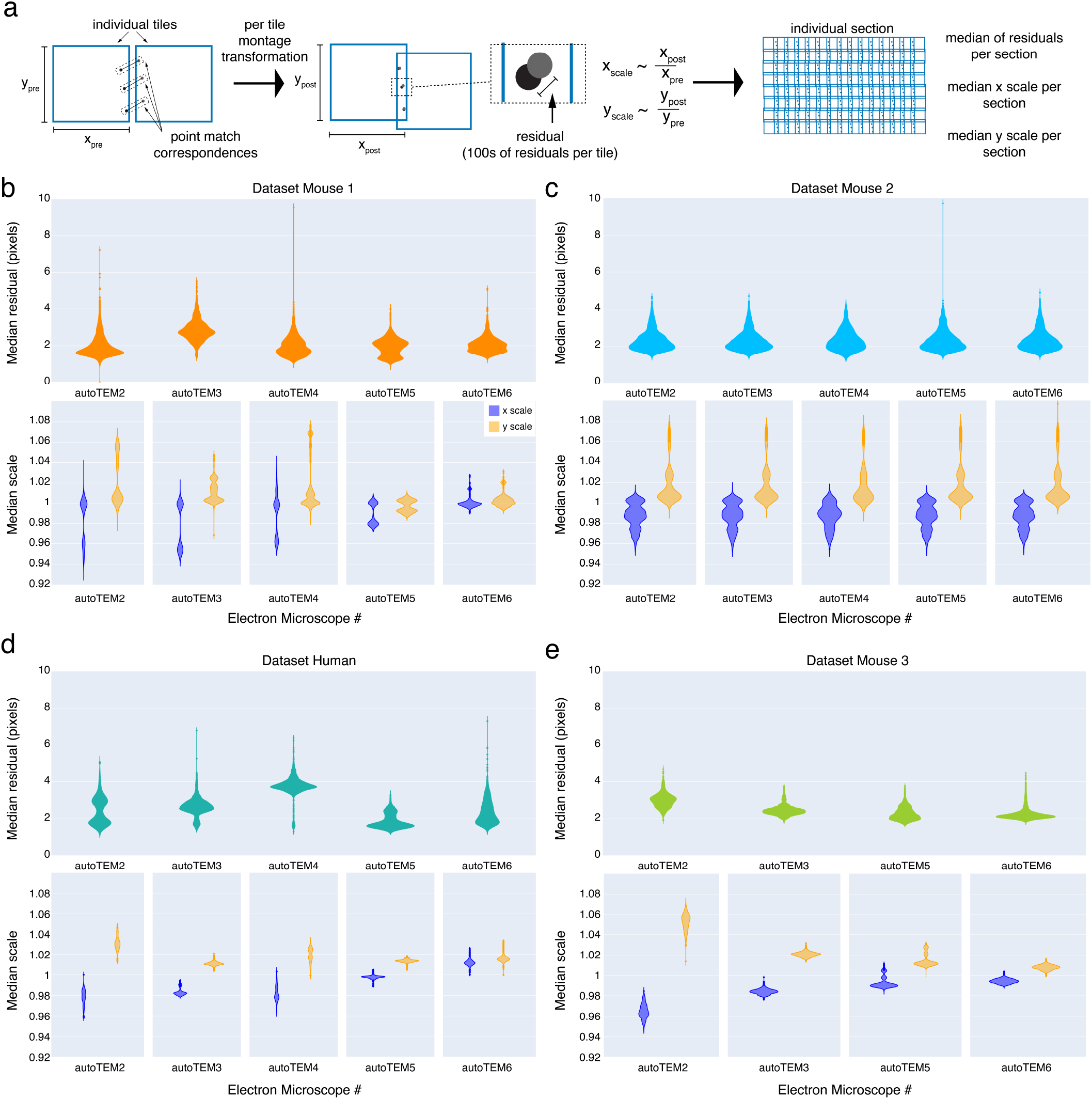
Performance of 2D stitching pipeline. (a) Schematic diagram explaining the computation of residuals between a pair of tile images. Panels b, c, d, and e show the median of tile residuals per section grouped by their acquisition TEMs (Top) and median residual distribution of *x* and *y* scales of the tile images (after montaging) per section grouped by their acquisition TEMs (Bottom) for all our datasets. (b) Mouse dataset 1. (c) Mouse dataset 2. (d) Human dataset. (e) Mouse dataset 3.

The global 3D alignment process produces a volume that is “roughly” aligned as the point correspondences are generated from montages materialized at 1% scale. This rough alignment provides a good initial approximation for fine alignment of the volume and for generating point correspondences at higher resolutions. The quality of global non-linear 3D alignment is measured by computing the angular residuals between pairs of sections (within a distance of 3 sections in *z*). The angular residual is computed using the point correspondences between a section and its neighbors. The angular residual is defined as the angle between two vectors formed by a point coordinate (from first section) and its corresponding point coordinate from a neighboring section. The origin of the two vectors is defined as the centroid of the first sections’ point coordinates. The median of the angular residuals is reported as a quality metric for the global 3D alignment for our datasets (see Fig. 6f). The quality metric ensures a high quality global non-linear 3D alignment of the sections in all three (*xy, yz, zx*) planes of the volume (see Fig. 6 for global non-linearly 3D aligned slices from mouse dataset 1 and supplemental figures 1, 2, and 3 for slices from other datasets).

**Figure 6:**
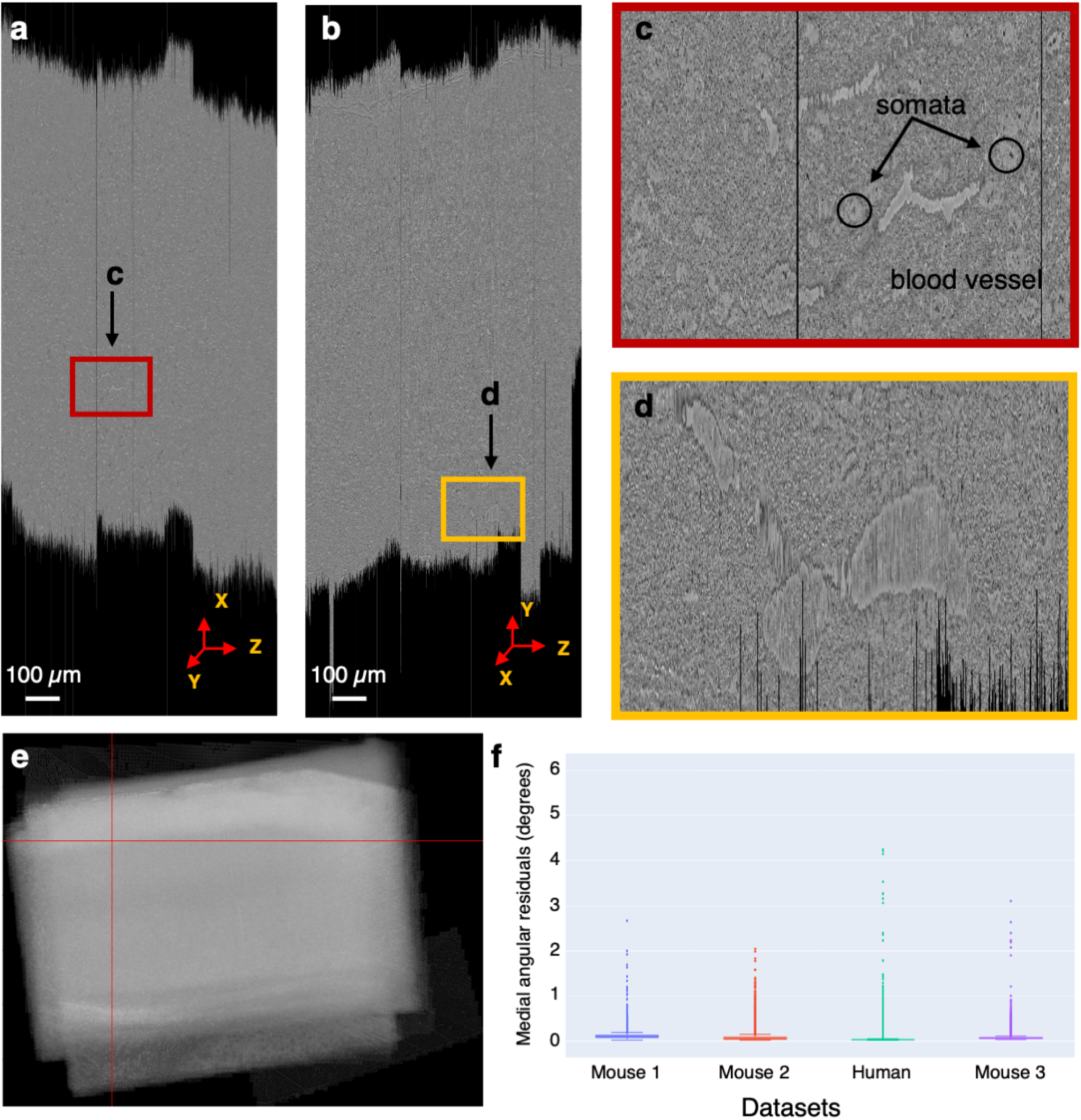
Global non-linear 3D aligned volume of the mouse dataset 1. a. View of the global non-linear 3D aligned volume from the *xz* plane. The figure shows the view of the global non-linear 3D alignment of the sections with the volume sliced at position marked by the red lines in (e). b. View of the global non-linear 3D aligned volume from the *yz* plane. Figure shows the view of the volume sliced at position marked by the red lines in (e). c. Zoomed-in area from (a) showing the quality of global non-linear 3D alignment in the *xz* plane. d. Zoomed-in area from (b) showing the quality of global non-linear 3D alignment in the *yz* plane. e. Maximum pixel intensity projection of the global non-linear 3D aligned sections in the z-axis showing the overall alignment of sections within the volume. The red lines represent the slicing location in both *xz* and *yz* plane for the cross section slices shown in (a) and (b). f. A plot showing the distribution of median angular residuals from serial sections grouped by the dataset.

Table 2 provides a comparison of both dataset acquisition times and their volume assembly. The acquisition times represent the serial sections imaged using 5 different ssTEMs running in parallel. Each of the dataset processing times are under the same infrastructure settings, but with several optimizations implemented in ASAP with every dataset. All of our datasets were processed in a time frame that matches or exceeds the acquisition time, thus achieving high-throughput volume assembly.

**Table 2:** Processing time comparison between acquisition system and ASAP. The acquisition times shown are based on serial sections imaged using 5 different ssTEMs running in parallel. The stitching time for all the datasets include the time it took to stitch all the serial sections including semi-automated QC and reprocessing sections that failed QC on the first run and the global 3D alignment. The stitching was done in a non-continuous fashion which included correctly uploading / re-uploading corrupted, duplicate sections, etc. Each section was stitched using a single node from the compute cluster. The different processing times of the different datasets reflect the optimization of the pipeline over time, while still keeping a throughput in pace with imaging acquisition.

| Dataset | # of sections | Acquisition time | ASAP processing time |
| --- | --- | --- | --- |
| Mouse dataset 1 | 26,500 | 6 months | 4 months |
| Mouse dataset 2 | 17,584 | 3 months | 6 weeks |
| Human dataset | 9,661 | 2 months | 4 weeks |
| Mouse dataset 3 | 17,309 | 3 months | 10 days |

### Application to other imaging pipelines: Array Tomography

The software infrastructure described in this manuscript can also be applied to fluorescence and multi-modal datasets such as array tomography (Figure 7). Array tomography presents some unique challenges for image processing because imaging can be performed in both light and electron microscopy. In addition, multiple channels can be imaged simultaneously and multiple rounds of imaging can be performed on the same physical sections with light microscopy [26]. To properly integrate all these images, in addition to the image processing steps of 2D stitching and alignment that apply to EM, the multiple rounds of light microscopy of the same section must be registered to one another, and the higher resolution EM data must be co-registered with the light microscopy data. Finally, alignments based on one set of images must be applied to the other rounds and/or modalities of data. The Render services allow for image processing steps to define new transformations on the image tiles without making copies of the data, including transformations that dramatically alter the scale of the images, such as when registering between EM and light microscopy data. The Render and point-match services provide a flexible framework for corresponding positions between tiles to be annotated, allowing those correspondences to be used as constraints in calculating the appropriate transformations at each step of the pipeline. The result is a highly multi-modal representation of the dataset that can be dynamically visualized in multiple channels and resolutions, at each step of the process through the integration of Render services with the Neuroglancer visualization tool (Figure 7).

**Figure 7:**
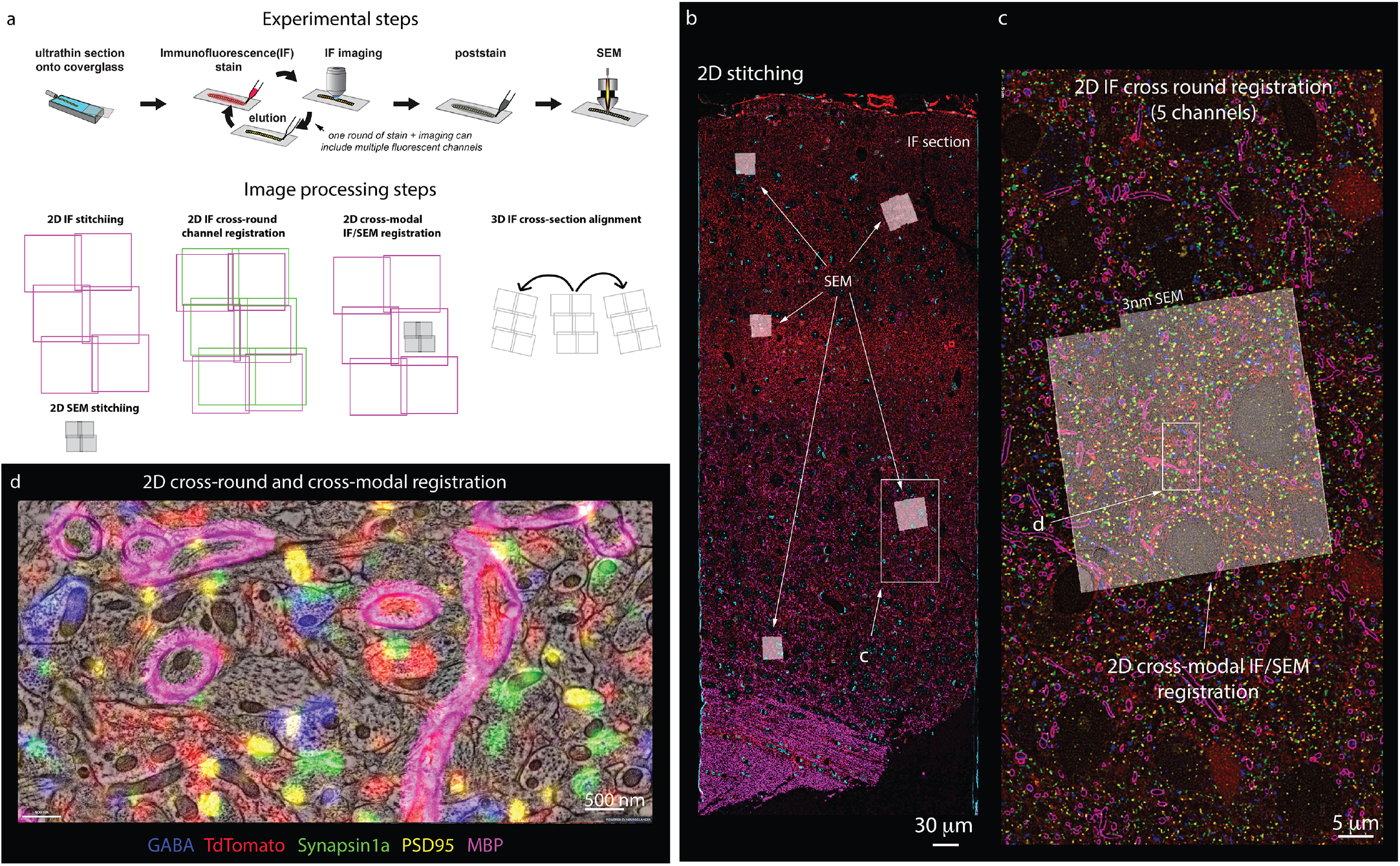
Stitching of multi-channel conjugate array tomography data. **a:Top)** Experimental steps in conjugate array tomography: Serial sections are collected onto glass coverslips, and exposed to multiple rounds of immuno-fluorescent (IF) staining, imaging and elution, followed by post-staining and imaging under a FESEM. **a:Bottom)** Schematic illustrating the substeps of image processing large scale conjugate array tomography data. 2D stitching must be performed on each round of IF imaging and EM imaging. Multiple rounds of IF imaging of the same physical section must be registered together to form a highly multiplexed IF image of that section. The higher resolution by typically smaller spatial scale FESEM data must then be registered to the lower resolution but larger spatial scale IF data for each individual 2D section and FESEM montage. Finally, alignments of the data across sections must be calculated from the IF, or alternatively EM datasets. In all cases, the transformations of each of these substeps must be composed to form a final coherent multi-modal, multi-resolution representation of the dataset. **b-d)** Screenshots of a processed dataset, rendered dynamically in Neuroglancer through the Render web services **b)**An overview image of a single section of conjugate array tomography data which shows the result of stitching and registering multiple rounds of IF an EM data. Channels shown are GABA (blue), TdTomato (Red), Synapsin1a (green), PSD95 (yellow), and MBP (purple). Small white box highlights the region shown in c. **c)** A zoom in of one area of the section where FESEM data was acquired, small white box shows the detailed region shown in d. **d)** A high resolution view of an area of FESEM data with IF data overlaid on top. One can observe the tight correspondence between the locations of IF signals and corresponding ultra-structural correlates, such a myelinated axons on MBP, and post-synaptic densities and PSD95.

**Figure 8:**
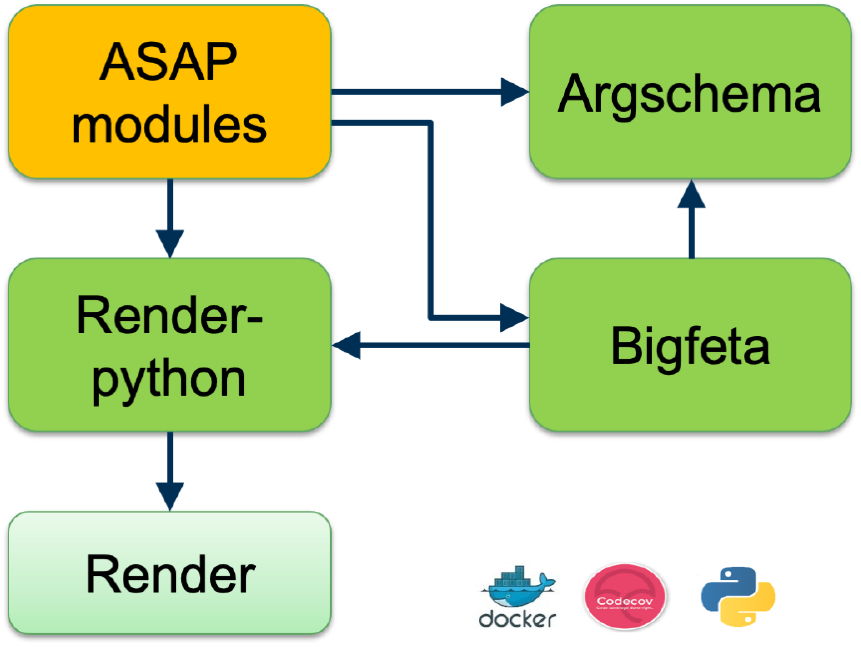
Set of software tools developed to perform peta-scale real-time stitching

## Discussion

The volume assembly pipeline ASAP was designed to produce a high throughput and high fidelity EM volume and is scalable, flexible, modular, upgradeable, and easily deployable on a variety of environments including large scale distributed systems. The pipeline leverages Render service’s capability of processing by means of meta-data operations and persisting data in databases. This largely facilitates multiple iterations of processing the data until a desired aligned volume is achieved. The need for rendering intermediate output is also eliminated at each iteration, since the output can be dynamically rendered by applying the meta-data associated with the images. This potentially saves computational time and resources in addition to increasing the throughput. Demonstrating its scalability, ASAP has been used to process several large scale datasets including a millimeter cube of mouse cortex that is already public in www.microns-explorer.org. Though ASAP is compatible with several strategies for fine alignment (Figure 1) the one used for all the datasets in this manuscript was SEAMLESS, which is described in [17].

The volume assembly pipeline maximizes the speed and quality of stitching and alignment on large scale datasets. One of the main improvements is the addition of a parameter optimization module that generates optimized sets of parameters for 2D stitching. This parameter optimization was introduced for montage solves in the mouse dataset 2, mouse dataset 3 and the human dataset. The use of optimization parameters resulted in less distorted montages with residuals within acceptable threshold values. It also compensated for some deviation in lens distortion correction accuracy, while reducing the number of iterations of processing.

### Quality assessment

In a software pipeline that processes tens of millions of images, it is essential to have automated metrics of quality control. The statistical metrics such as median absolute deviation (MAD) of the image scales to auto detect deformed montages combined with detecting other stitching issues by the QC module facilitates faster processing while ensuring that the stitched sections meet the QC criteria. Also, early detection of poor point correspondences by the QC module drastically reduces the need for reprocessing montages through several iterations. About 2% of sections undergo this re-computation of point correspondences at a higher scale. Speed-up is also achieved by automating data transfer and ingestion into our volume assembly workflow from imaging. This is achieved by means of automatically querying the imaging database for sections that have been imaged and have passed imaging QC [1]. The meta-data of the QC passed sections are automatically ingested into the volume assembly workflow, which also triggers the stitching process. The imaging database was not developed during imaging of mouse dataset 1, hence the status of imaging and QC for each section was maintained in a spreadsheet and manually updated.

ASAP is capable of handling re-imaged serial sections without overwriting the meta-data for its earlier versions during processing. Also, the system is capable of handling missing sections (in case of serial section loss during sectioning or aperture burst/damage during imaging) and partial sections (sections that are cut partially from the volume). The missing sections are treated as “gaps” in the volume and have minimal impact on the quality of alignment. Currently, the pipeline has successfully handled a gap of 3 consecutive sections (and 5 consecutive sections for the human dataset) in the volume. Feature-based computation of point correspondences is effective in finding features across sections with gaps between them and also robust to contrast and scale variations between image pairs.

The software stack includes capabilities to interface with different solvers through BigFeta including a scipy.sparse-based solver and the interfaces provided by PETSc ([22], [23], [24]). This has allowed us to non-linearly globally 3D align an entire volume on a single workstation as well as on a distributed system. Our code-base was also improved to allow for re-processing individual sections that are re-imaged and inserting them in existing global non-linear 3D aligned volume. In addition to file storage, our software tools now support object stores using an S3 API such as Ceph, Cloudian, and AWS, enabling real-time processing of large-scale datasets in the cloud as well as on-premises. The entire software stack is designed and developed using open-source dependencies and licensed under the permissive Allen Institute Software License. Also, our software stack and its modules are containerized allowing rapid deployment and portability. It also includes integration tests for each module for seamless development and code coverage. Automated processing of EM datasets can be accomplished with a custom workflow based on an open-source workflow manager (BlueSky) that is well suited to incorporate complex workflows with readable, flexibility workflow diagrams allowing rapid development.

### Image processing at the speed of imaging

The reconstruction of neural circuits requires high spatial resolution images provided by EM and drastic advances made in the field of EM connectomics ([1, 25, 31]) make it suitable for imaging large scale EM volumes and produce dense reconstructions. ASAP aligns well with such large scale EM volume production systems facilitating seamless processing of data through automated data ingestion, 2D stitching, 3D alignment, and QC - all chained together as a continuous process. Developing a pipeline that can produce a high throughput, high fidelity EM volume at a rate better than imaging was the most challenging problem. In addition, we invested heavily to develop a set of software tools that is modular, easily adaptable and upgradeable to new algorithms, computing systems and other domains, and able to run in a production level setting.

The offline processing duration of all our datasets using ASAP have shown to exceed the speed of imaging. ASAP is capable of processing the datasets in parallel with the imaging session with sufficient computational resources. The mouse dataset 1 was processed in parallel with imaging (stitching of serial sections) followed by a chunk based global 3D alignment (first iteration). Efficient data transfer from the multi-scope infrastructure coupled with automated processing capabilities of ASAP, assisted in processing of the mouse dataset 1 in parallel to imaging and at speeds that match the imaging. The *em stitch* software package leverages the GPU-based computations on the scope for imaging QC to stitch the montages on-scope. This accelerates the stitching process and the rapid feedback loop between imaging and volume assembly. Though our current processing rate already out-performs image acquisition, the next step is to perform the image processing in real time, ideally close to the microscopes and as images are collected. Such strategy has been proposed by Jeff Lichtman and colleagues ([30]) and there are many aspects of the work presented here that will facilitate transition to on-scope real time stitching and alignment.

### Scaling to larger volumes and across image modalities

Our pipeline was developed with a focus on standardization and was built entirely with open source libraries as an open source package. Our intention is for others to use and potentially scale it beyond the work described in this manuscript. As we demonstrate in figure 7, the use of ASAP goes well beyond electron microscopy, and it is being used in fluorescent data as well. The modularity of ASAP can be leveraged to include GPU based algorithms at various stages of the pipeline, thus paving the way for further increase in throughput. Processing in parallel with imaging, we were able to stitch and globally non-linearly 3D align 2 PB of EM images from the 1 mm^3^ mouse visual cortex at synapse resolution within a period of ~4 months, and other peta-scale datasets with a montaging rate exceeding the imaging rate. With improvements made to the pipeline, stitching and global non-linear 3D alignment of a dataset similar in size took just 10 days of processing time for mouse dataset 3. This throughput makes the volume assembly pipeline suitable for processing exascale datasets that spans larger cortical areas of the brain across species. Although the pipeline was designed for EM connectomics, it can be easily adapted to process datasets from any other domain of image sets that share the basic underlying assumptions in imaging.

## Methods

### Imaging with Electron Microscopy

Three of the samples processed by the infrastructure described in this manuscript originated from mice. All procedures were carried out in accordance with Institutional Animal Care and Use Committee approval at the Allen Institute for Brain Science. All mice were housed in individually ventilated cages, 20–26°C, 30–70% Relative Humidity, with a 12 h light/dark cycle. Mice(CamK2a-tTA/CamK2-Cre/Ai93, CamKII-tTA/tetO-GCaMP6s, Slc-Cre/GCaMP6s). Preparation of samples was performed as described earlier ([1]), briefly mice were transcardially perfused with a fixative mixture of 2.5% para-formaldehyde and 1.25% glutaraldehyde in buffer. After dissection, slices were cut with a vibratome and post-fixed for 12–48 h. Human surgical specimen was obtained from local hospital in collaboration with local neurosurgeon. The sample collection was approved by the Western Institutional Review Board (Protocol # SNI 0405). Patient provided informed consent and experimental procedures were approved by hospital institute review boards before commencing the study. A block of tissue approximately 1 × 1 × 1 cm of anteromedial temporal lobe was obtained from a patient undergoing acute surgical treatment for epilepsy. This sample was excised in the process of accessing the underlying epileptic focus. Immediately after excision, the sample was placed into a fixative solution of 2.5% paraformaldehyde, 1.25% glutaraldehyde, 2mM calcium chloride, in 0.08 M sodium cacodylate buffer for 72 h. Samples were then trimmed and sectioned with a vibratome to 1000 μm thick slices, and placed back in fixative for ~96 h. After fixation, slices of mouse and human were extensively washed and prepared for reduced osmium treatment (rOTO) based on the protocol of Hua et al. [27] potassium ferricyanide was used to reduce Osmium tetroxide and thiocarbohydrazide (TCH) for further intensification of the staining. Uranyl acetate and Lead aspartate were used to enhance contrast. After resin embedding, ultrathin sections (40 nm or 45 nm) were manually cut in a Leica UC7 ultra-microtome and a RMC Atumtome. After sectioning, the samples were loaded into the automated Transmission Electron Microscopes (autoTEM) and we followed the TEM operation routine (described in [1] and [25]) to bring up the HT voltage and filament current and then align the beam. Calibration of the autoTEM involved tape and tension calibration for bar-code reading, measuring beam rotation and camera pixels, and stage alignment. After which, EM imaging was started. The mouse datasets were obtained from primary visual cortex and higher visual areas, the human dataset was obtained from the Medial Temporal Gyrus (MTG).

### Image Catcher (Aloha) service

Aloha is a core component of our acquisition infrastructure designed to facilitate the transfer and pre-processing of images intended for the image processing workflow. Aloha is implemented as a scale-out python web service using flask/gunicorn. This service is designed to accept image arrays defined by a flat-buffers protocol and atomically write them in a designated location in losslessly-compressed tiff format. While the array is in memory, the service also writes progressively downsampled versions of that image (MIPmaps) to another designated location. By using the uri-handler library [16], aloha can write to various cloud providers and on-premises object storage systems as well as file system-based storage. The Aloha library includes a set of client scripts which allow uploading from an existing autoTEM-defined directory as well as utilities to encode numpy arrays for the REST API. Aloha web service is configured to interact with the piTEAM’s TEMdb backend and tracks the state of transfers in a set of custom fields. In the automated workflow, a process queries these fields in order to ingest completed montage sets to the volume assembly workflow. Aloha can be easily replaced with a data transfer module of choice based on the imaging infrastructure and the volume assembly workflow allowing for modularity.

### Render services

The Render services are a core component of the infrastructure. They provide the main logic for image transformation, interpolation and rendering. They also provide a rich API:

- A REST API for creating and manipulating collections of tiles or image “boxes” (also called canvases; canvases are regions that can span multiple and partial tiles).
- A REST API for accessing image tile, section and stack meta information: For example the number of tiles, dimensions, ids, camera setup.
- A REST API and core logic for rendering/materializing image tiles/canvases, arbitrary regions that span a number of (or partial) tiles, or even whole sections. In that capacity, it is used to reflect the current state of any given tile collection. (This is invaluable to proofreading intermediate stitching results). In combination with dynamic rendering (i.e. rendering that is not based on materializing image files to storage), the Render provides light-weight feedback to detect imaging and stitching issues.

The Render services are backed by a MongoDB document store that contains all tile/canvas data including tile transformations. Both the Render services and the MongoDB document store are supported by dedicated hardware. The Render services code base is available and documented here: https://github.com/saalfeldlab/render

### Point-match service

A time-consuming and CPU-intensive process in the volume assembly pipeline is the computation of point correspondences between image tile pairs since this is the only stage of processing where the image data is read in memory besides the process of rendering the aligned volume to disk. Persisting this data is therefore invaluable. Robust rotation and contrast invariant correspondence candidates are generated using SIFT [19]. These candidates are then filtered by their consensus with respect to an optimal geometric transformation, in our case an affine transformation. We use a local optimization variant of RANSAC [21] followed by robust regression [5]. Local optimization means that, instead of picking the ‘winner’ from a minimal set of candidates as in classic RANSAC, we iteratively optimize the transformation using all inlier candidates and then update the inlier set. The ‘winner’ of this procedure (the largest set of inliers) is then further trimmed by iteratively removing candidates with a residual larger then 3 standard deviations of the residual distribution with respect to the optimal transformation and then re-optimizing the transformation. We use direct least-squares fits to optimize transformations. The computed point correspondences are stored in a database and can be retrieved/modified using the point-match service. The advantage of such a database is that it is agnostic to the source of point correspondences. Therefore, it can receive input from the point-match generator, regardless of the method of point-match generation such as SURF, SIFT, phase-correlation, etc.

### Render-python API

The other core component of the software stack include *render-python*, a Python API client and transformation library that interacts with both asap-modules and the Render services. The render-python components interact with Render service Java clients that perform computationally expensive operations locally to avoid taxing Render services running on centralized shared hardware.

*render-python* is a python-based API client and transformation library that replicates the data models in the Render services. While Render services utilize the mpicbg Fiji library to implement transformations, renderpython reproduces these using using numpy to enable analysis in a python ecosystem. render-python is continuously integration tested against Render for compatibility and provides dynamic access to the database and client scripts provided by Render.

Besides render-python, ASAP interfaces with other tools for solving for transformations and for visualizations. A description of these tools are as follows;

#### BigFeta

The *BigFeta* package implements a python-based sparse solver implementation of alignment problems based on the formulation in EMAligner [12]. In addition to optimizations and new transform functionality, BigFeta extends the previous approach to use PETSc (petsc.org) for scalable computation and allows input and output using render-python objects as well as JSON file, MongoDB document store, and Render services interaction.

#### em stitch

*em stitch* includes tools based on BigFeta and render-python for use as a standalone montage processing package without connecting to a database or REST API, ideal for online processing utilizing the same hardware running a microscope. Importantly, *em stitch* includes a module to derive a mesh-based lens distortion correction from highly-overlapping calibration acquisitions and pre-generated point correspondences.

#### vizrelay

*vizrelay* is a configurable micro-service designed to build links from a Render services instance to a Neuroglancer-based service and issue redirects to facilitate visualization. This provides a useful mechanism for setting Neuroglancer defaults, such as the extent of the volume or color channel options when reviewing alignments.

### ASAP modules

The ASAP volume assembly pipeline includes a series of modules developed using Python and the renderpython library that implement work-flow tasks with standardized input and output formatting. The submodules in ASAP include scripts to execute a series of tasks at each stage of the volume assembly pipeline. Some of the workflow tasks included in ASAP are as follows:

- *asap.dataimport:* import image (tile) meta-data to the Render services from custom microscope files, generate MIPmaps and update the meta-data, render downsampled version of the montaged serial section.
- *asap.mesh_lens_correction*: include scripts to compute the lens distortion correction transformation.
- *asap.pointmatch*: generate tile pairs (see Fig. 1d) and point-correspondences for stitching and alignment.
- *asap.point_match_optimization*: Find the best possible set of parameters for a given set of tile pairs.
- *asap.solver*: interface with BigFeta solver for stitching the serial sections.
- *asap.em_montage_qc*: generate QC statistics on the stitched sections as explained in Section Automated petascale stitching.
- *asap.rough_align*: compute per-section transformation for 3D alignment, and scale them to match their original layered montage collection and generate new meta-data describing the alignment at full resolution.
- *asap.register*: register an individual section with another section in a chunk. This module is typically used to align re-imaged sections to an already aligned volume.
- *asap.materialize*: materialize final volume as well as downsampled versions of sections in a variety of supported formats.

ASAP modules are schematized for ease of use with *argschema*, an extension of the marshmallow python package which allows marshalling of command-line arguments and input files.

### Montage Parameter Optimization

BigFeta implements the optimization described by Khairy et al. [12] such that the regularization parameter *λ* can differently constrain distinct terms of a transformation such as the translation, affine, and polynomial factors on an individual tile basis.

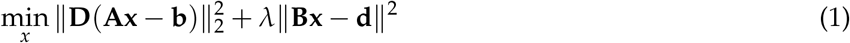

Montage quality in ASAP is evaluated by metrics of residuals and rigidity of the output montage (Figure 3, Figure 4). For tile deformations that are well-defined by an affine transformation, these metrics are most impacted by the translation and affine regularization parameters (*λ*) used in the BigFeta solver step (equation 1). As the optimal configuration of these values can be impacted by the accuracy of the initialization as well as the weight and distribution of point correspondences, it is sometimes necessary to re-evaluate the regularization parameters for different imaging, tissue, or pre-processing conditions. We provide an “optimization” module, asap.solver.montage_optimization, which leverages the fast solving capabilities of BigFeta to sweep across a range of regularization parameters and provide an acceptable set of parameters given targets for correspondence residual in pixels and tile scale MAD value.

### 3D Realignment

A common use case after 3D global alignment involves re-aligning a subset of the dataset while maintaining the global alignment reached for the rest of the volume. The asap.solver.realign_zs module implements this operation by increasing the *λ* parameters in equation 1 for neighboring sections while allowing custom inputs for the sections that need to be realigned. As such, it is possible to integrate re-montaged sections, recomputed point correspondences, or more deformable transformations into an existing 3D alignment without requiring changes on the global scale. For all the datasets presented in this manuscript, after global alignment the data was then transferred for fine alignment using SEAMLESS [17]). The fine alignment was performed by the team of Sebastian Seung in Princeton or ZettaAI.

### Chunk Fusion

The asap.fusion package provides modules to support chunk-based 3D alignment workflows. The 3D aligned chunks can then be fused together. asap.fusion.register_adjacent_stack provides utilities to register overlapping 3D aligned chunks using translation, rigid, similarity, or affine transformations. Then, given a JSON-defined tree describing the layout and relative transformation between chunks, asap.fusion.fuse_stacks will assemble meta-data representing a combined volume using Render’s “InterpolatedTransform” to interpolate between independently optimized transformations in the overlap region of two chunks.

### BlueSky workflow engine for automated processing

The automated workflow engine called BlueSky was developed in Django backed by a PostgreSQL database with stable backups, graceful restarts, and easy migrations. It provides a web based user interface for the user to visualize, run, edit running jobs at various stages in the workflow. BlueSky uses Celery and RabbitMQ to run workflow tasks in diverse computing environments, from local execution on a workstation to remote execution using a compute cluster (PBS, MOAB, SLURM). BlueSky is flexible in terms of designing complex workflows as the workflow diagrams (see supplementary Figs. 4 and 5) can be specified in readable formats such as YAML, JSON or Django allowing rapid development. BlueSky can be used for many different purposes, but for the image processing task related to this manuscript the workflow includes the following steps:

(1) ingest montage sets, (2) generate MIPmaps, (3) apply MIPmaps, (4) wait for the assigned lens correction transform, (5) apply the lens correction transform, (6) extract tile pairs for determining point correspondences, (7) generate 2D montage point correspondences, (8) run the 2D montage solver, (9) automatically check for defects, (10) place potential defects in a manual QC queue and (11) generate downsampled montage. BlueSky is publicly available at GitHub (https://github.com/AllenInstitute/blue_sky_workflow_engine). The volume assembly workflow is designed to use BlueSky workflow engine for processing our datasets. The custom EM volume assembly workflow (https://github.com/AllenInstitute/em_imaging_workflow) facilitates continuous processing of the datasets at speeds that match or exceed data acquisition rates (see supplementary Figs. 6 and 7).

## Acknowledgements

We thank our project manager Shelby Suckow for her exceptional work on keeping us aligned and on time. We thank Allan Jones and Gerry Rubin for starting the conversation that lead to this collaboration and for their support and leadership. We also thank Hongkui Zeng and Christof Koch for their support and leadership. We thank D. Brittain, M. Scott and J. Borseth for their help collecting and imaging the EM datasets. We thank Sebastian Seung, Thomas Macrina, Nico Kemnitz, Manuel Castro, Dodam Ih, and Sergiy Popovych in Princeton University and ZettaAI for discussions and feedback on image processing strategies and improvements. We thank Brian Youngstrom, Stuart Kendrick and the Allen Institute IT team for support with infrastructure, data management and data transfer. We thank Jay Borseth, DeepZoom LLC for his contributions to *em stitch*. We thank Andreas Tolias, Jacob Reimer and their teams at the Baylor College of Medicine for providing mice used for electron microscopy. We also thank Saskia de Vries, Jerome Lecoq, Jack Waters and their teams at the Allen Institute for providing mice used for electron microscopy. This work was supported by the Intelligence Advanced Research Projects Activity (IARPA) of the Department of Interior/Interior Business Center (DoI/IBC) through contract number D16PC00004; and by Allen Institute for Brain Science. The views and conclusions contained herein are those of the authors and should not be interpreted as representing the official policies or endorsements, either expressed or implied, of the funding sources including IARPA, DoI/IBC, or the U.S. Government. The authors wish to thank the Allen Institute founder, Paul G. Allen, for his vision, encouragement, and support.

## Author Contributions

N.M.C. and R. C. Reid conceptualized the project and funding acquisition. G.M., R.T. validated and optimized the pipeline software (asap-modules) and executed the stitched and aligning of the datasets. D.K. developed EM aligner. T. F. developed BlueSky workflow manager and the volume assembly workflow. Saalfeld, S. and E.T.T are primary developers of the Render services. K.K. is the primary developer of the MATLAB version of EM aligner. R.T. developed Aloha with help of S.K., K. K. conceptualized the stitching pipeline using Render services. E. P. developed multi-channel support for asap-modules and vizrelay. G.M., R. T, F. C., Seshamani, S. and E.P. developed the software packages (asap-modules, render-python) and maintained the infrastructure and continuous integration testing used by the Render backed pipeline. J.B., D.B, N.M.C, M.T and W.Y. contributed to the generation of mouse EM data. J.B., D.B, N.M.C, E.L., J.N., M.T and W.Y. contributed to the generation of human EM data. Seshamani, S. and F.C. generated Array tomography data. G.M., R.T. and N.M.C. wrote the manuscript with contributions from other authors.

