## Supplemental Material for "A Scalable and Modular Automated Pipeline for Stitching of Large Electron Microscopy Datasets" (from another version of this preprint: version 1, 2021-11-25)

immediate

November 25, 2021

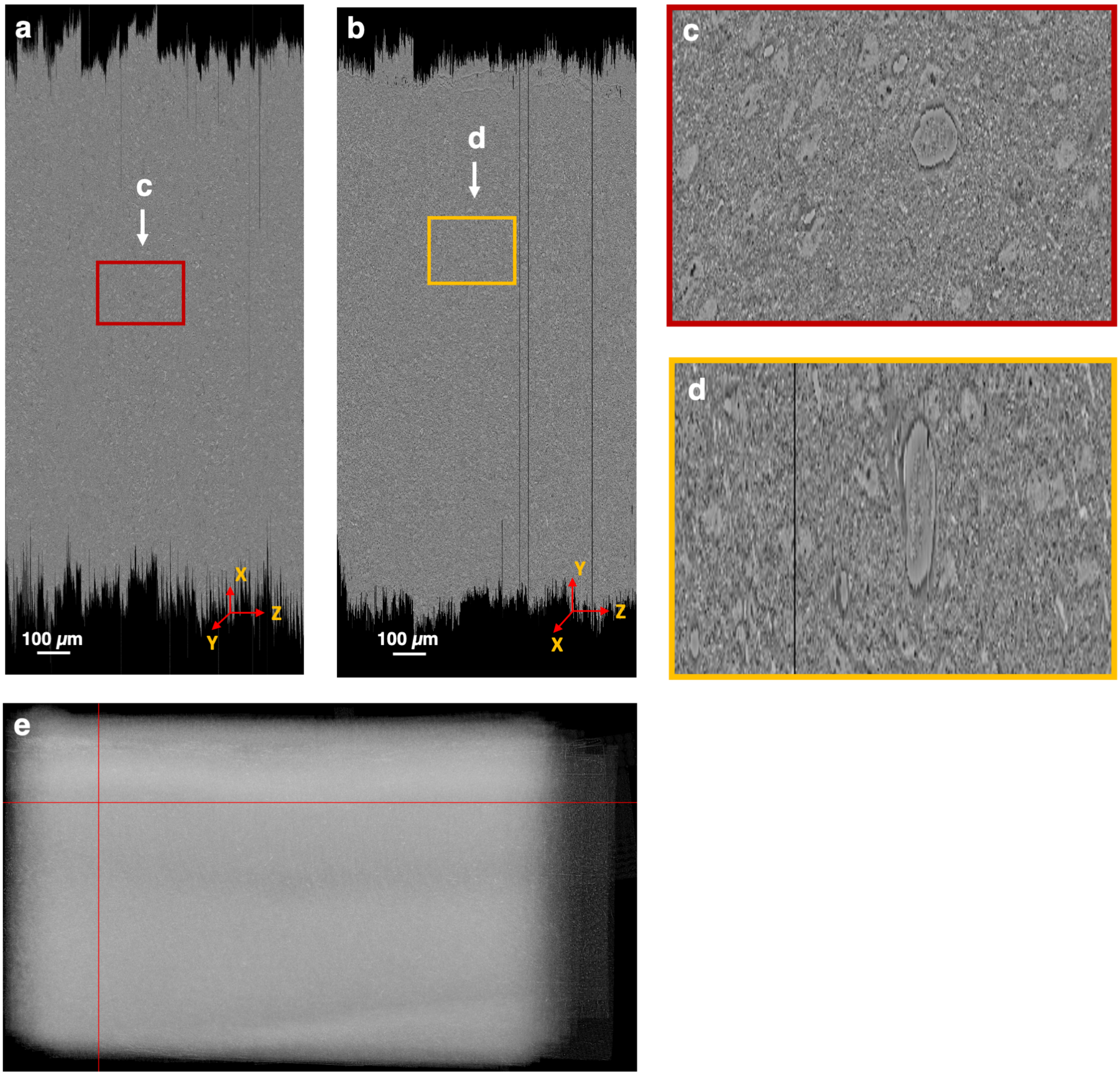


Figure 1: **Rough aligned volume of the mouse dataset 2**. a. View of the rough aligned volume from the xz plane. The figure shows the view of the rough alignment of the sections with the volume sliced at position marked by the red lines in (e). b. View of the rough aligned volume from the yz plane. Figure shows the view of the volume sliced at position marked by the red lines in (e). c. Zoomed-in area from (a) showing the quality of rough alignment in the xz plane. d. Zoomed-in area from (b) showing the quality of rough alignment in the yz plane. e. Maximum pixel intensity projection of the rough aligned sections in the z-axis showing the overall alignment of sections within the volume. The red lines represent the slicing location in both xz and yz plane for the cross section figures in (a) and (b).

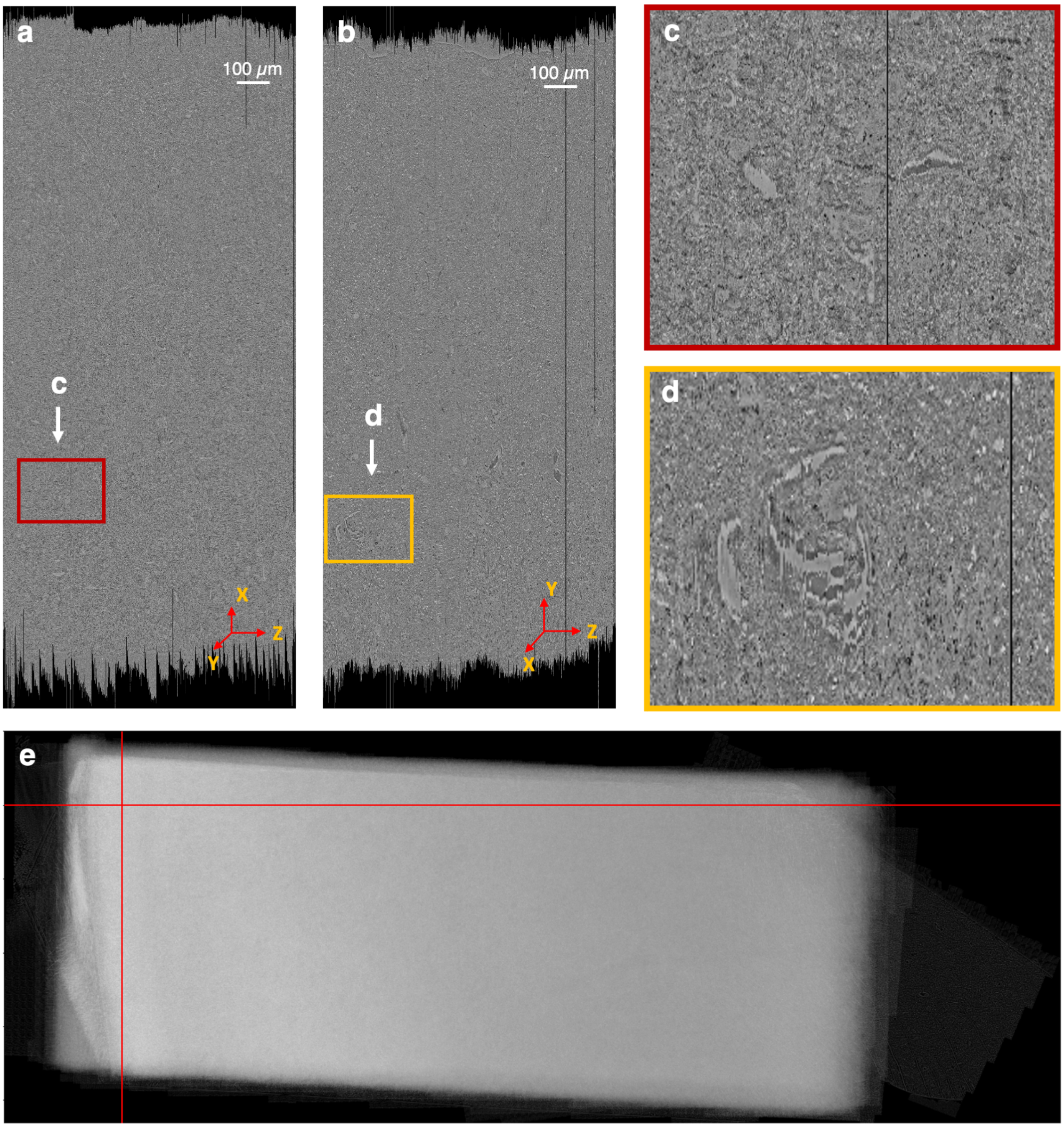


Figure 2: **Rough aligned volume of the human dataset**. a. View of the rough aligned volume from the xz plane. The figure shows the view of the rough alignment of the sections with the volume sliced at position marked by the red lines in (e). b. View of the rough aligned volume from the yz plane. Figure shows the view of the volume sliced at position marked by the red lines in (e). c. Zoomed-in area from (a) showing the quality of rough alignment in the xz plane. d. Zoomed-in area from (b) showing the quality of rough alignment in the yz plane. e. Maximum pixel intensity projection of the rough aligned sections in the z-axis showing the overall alignment of sections within the volume. The red lines represent the slicing location in both xz and yz plane for the cross section figures in (a) and (b).

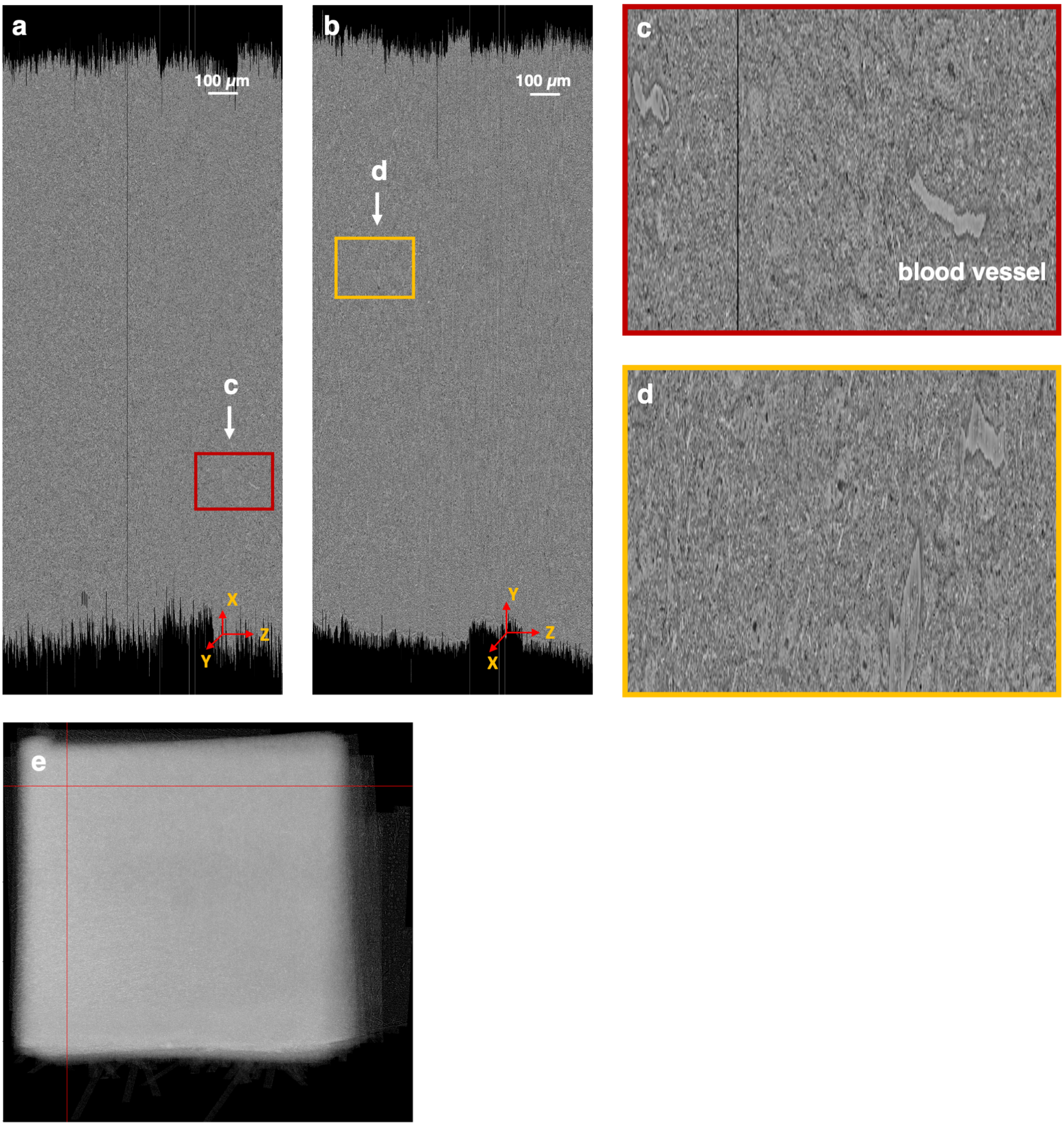


Figure 3: **Rough aligned volume of the mouse dataset 3**. a. View of the rough aligned volume from the xz plane. The figure shows the view of the rough alignment of the sections with the volume sliced at position marked by the red lines in (e). b. View of the rough aligned volume from the yz plane. Figure shows the view of the volume sliced at position marked by the red lines in (e). c. Zoomed-in area from (a) showing the quality of rough alignment in the xz plane. d. Zoomed-in area from (b) showing the quality of rough alignment in the yz plane. e. Maximum pixel intensity projection of the rough aligned sections in the z-axis showing the overall alignment of sections within the volume. The red lines represent the slicing location in both xz and yz plane for the cross section figures in (a) and (b).

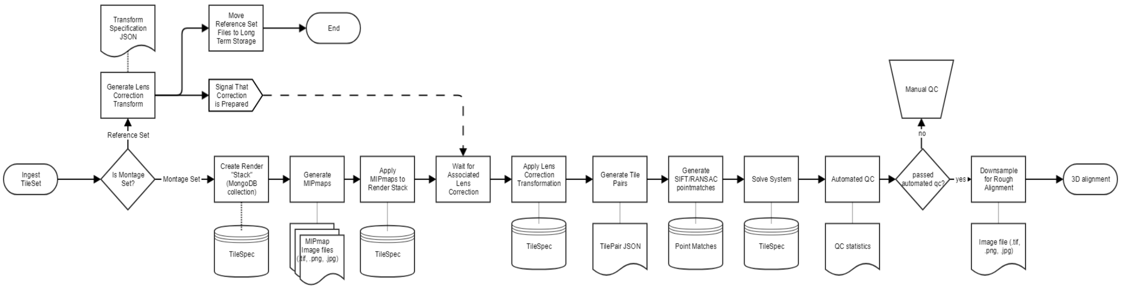


Figure 4: IARPA MICrONS Phase 2 Montage workflow with support for Lens Correction and Manual Intervention

```
graph:
    - [ "ingest_tile_sets", [ "generate_lens_correction", "generate_em_montage" ] ]
    - [ "generate_lens_correction", [ "move_reference_set" ] ]
    - [ "generate_em_montage", [ "generate_mipmaps" ] ]
    - [ "generate_mipmaps", [ "apply_mipmaps" ] ]
    - [ "apply_mipmaps", [ "wait_for_lens_correction" ] ]
    - [ "move_reference_set", [ "wait_for_lens_correction" ] ]
    - [ "wait_for_lens_correction", [ "apply_lens_correction_new" ] ]
    - [ "apply_lens_correction_new", [ "create_tile_pairs" ] ]
    - [ "create_tile_pairs", [ "2d_montage_point_match" ] ]
    - [ "2d_montage_point_match", [ "solver" ] ]
    - [ "solver", [ "detect_defects" ] ]
    - [ "detect_defects", [ "wait_for_manual_qc" ] ]
    - [ "wait_for_manual_qc", [ "downsample", "move_raw_montage_set" ] ]
    - [ "downsample", [ "alignment" ] ]
```

Figure 5: EM workflow diagram from 4 represented in YAML as DAG

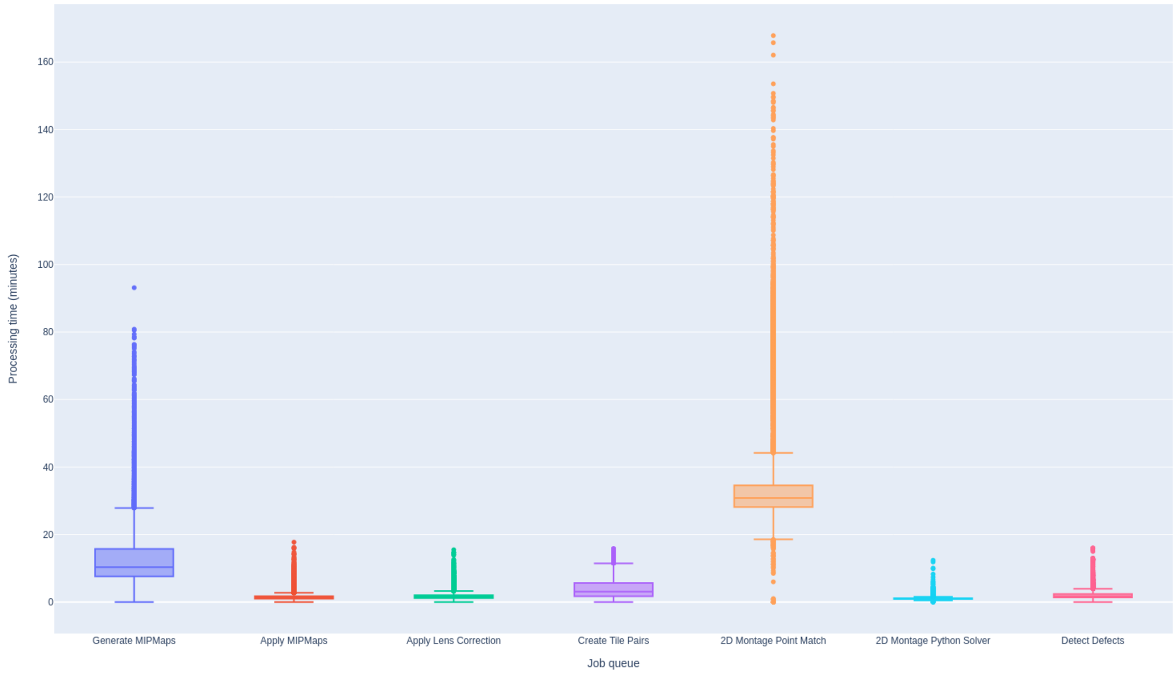


Figure 6: **Performance of BlueSky workflow manager**. Time spent by the sections from all the datasets in each job queue in the montaging workflow. The processing times include the duration between the time the job started running in a node and the time the node releases the job as successfully completed. Processing times shown are based on running the job in a single computing node in every job queue.

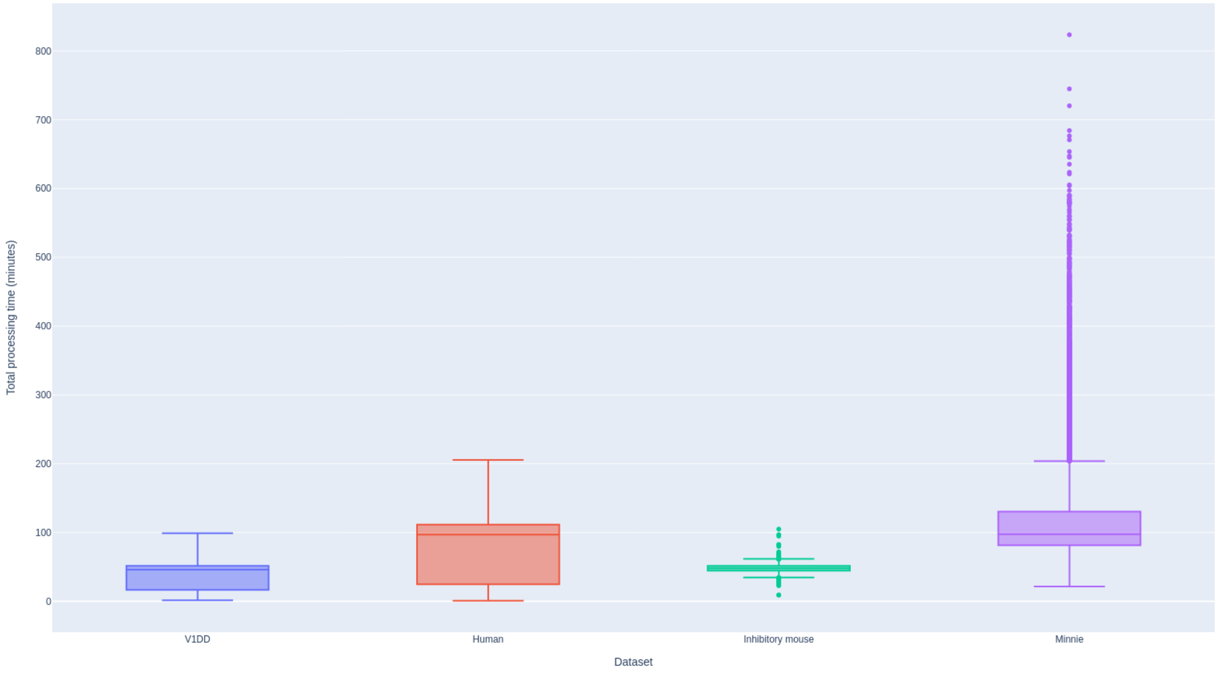


Figure 7: **Performance of BlueSky workflow manager**. Total processing time for sections montaged using the BlueSky workflow manager for all the datasets. Processing times shown are based on running the job in a single computing node in every job queue.
